# Previous estradiol treatment during midlife maintains transcriptional regulation of memory-related proteins by ERα in the hippocampus in a rat model of menopause

**DOI:** 10.1101/2021.02.23.432513

**Authors:** Nina E. Baumgartner, Katelyn L. Black, Shannon M. McQuillen, Jill M. Daniel

## Abstract

Previous midlife estradiol treatment, like continuous treatment, improves memory and results in lasting increases in hippocampal levels of estrogen receptor (ER) α and ER-dependent transcription in ovariectomized rodents. We hypothesized that previous and continuous midlife estradiol act to specifically increase levels of nuclear ERα, resulting in transcriptional regulation of proteins that mediate estrogen effects on memory. Ovariectomized middle-aged rats received estradiol or vehicle capsule implants. After 40 days, rats initially receiving vehicle received another vehicle capsule (Vehicle). Rats initially receiving estradiol received either another estradiol (Continuous Estradiol) or a vehicle (Previous Estradiol) capsule. One month later, hippocampal genes and proteins were analyzed. Continuous and previous estradiol increased levels of nuclear, but not membrane or cytosolic ERα and had no effect on *Esr1*. Continuous and previous estradiol impacted gene expression and/or protein levels of mediators of estrogenic action on memory including ChAT, BDNF, and PSD-95. Findings demonstrate a long-lasting role for hippocampal ERα as a transcriptional regulator of memory following termination of previous estradiol treatment in a rat model of menopause.

## 1. Introduction

Decades of research support the idea that estrogens play an important role in modulating memory in the aging female brain (Koebele and Bimonte-Nelson, 2017; Luine and Frankfurt, 2020). Declines in ovarian hormones following menopause coincide with increased risk of pathological and non-pathological cognitive decline. Estrogens administered near the onset of loss of ovarian function have been shown to improve cognition in humans and rodents and enhance function of the hippocampus, a brain region crucial for memory (Daniel et al., 2015; Maki et al., 2011). However, due to serious health risks of long-term estrogen use (Chen et al., 2006), current guidelines recommend that individuals who do choose to use estrogens to treat menopause symptoms do so for only a few years near menopause (Santen et al., 2010).

In an aging ovariectomized rodent model of menopause, our lab has shown that short-term estrogen use during midlife has lasting effects on the brain and cognition. Forty days of estradiol exposure immediately following midlife ovariectomy resulted in enhanced performance on the hippocampal-dependent radial arm maze up to seven months after estradiol treatment had been ended, effects that were comparable to ongoing estradiol treatment (Rodgers et al., 2010). This initial finding in our lab demonstrated that estrogens administered for a short-term period immediately after the loss of ovarian function can have lasting benefits for memory similar to those exerted by ongoing estradiol exposure. Since then, we have replicated this lasting impact of previous exposure to midlife estradiol on memory in rats (Black et al., 2018; Black et al., 2016; Witty et al., 2013) and in mice (Pollard et al., 2018). Evidence in nonhuman primates receiving short-term estrogen use in midlife shows similar results. Ovariectomized rhesus monkeys that received 11 months of cyclic estradiol injections displayed enhanced performance on a memory task one year after termination of hormone treatment (Baxter et al., 2018). Finally, in women who undergo surgical menopause (oophorectomy) earlier in life than natural menopause, estrogen treatment until the time at which natural menopause would occur is recommended and is associated with decreased risk of cognitive impairment later in life (Bove et al., 2014; Rocca et al., 2014). Collectively, these studies demonstrate across multiple species that exposure to estrogens immediately following loss of ovarian function can improve cognition long after hormone treatment has ended.

The ability of previous midlife estradiol exposure to enhance cognitive aging long-term is related to its ability to impact levels of estrogen receptors in the hippocampus. Estrogens exert their effects on the brain by acting on estrogen receptors, including the classic nuclear steroid receptor estrogen receptors (ER) α. Previous midlife estradiol exposure resulted in lasting increases in hippocampal levels of ERα eight months after termination of estradiol treatment (Rodgers et al., 2010). Subsequent work in our lab and others suggest a causal relationship between increased hippocampal ERα and memory. For instance, increasing hippocampal ERα using lenti-viral vectors enhances performance on the radial arm maze in aged ovariectomized rats (Witty et al., 2012). Additionally, pharmacologically antagonizing ERα using ICI 182780 prevents the memory benefits shown previously with our short-term estradiol model in aged ovariectomized rats (Black et al., 2016). Finally, certain polymorphisms in *Esr1,* the gene that encodes for ERα, may impact cognitive function (Ma et al., 2014; Yaffe et al., 2009) and increase risk of Alzheimer’s disease in postmenopausal women (Ma et al., 2009; Ryan et al., 2014), demonstrating that ERα can impact memory long after cessation of ovarian function.

Sustained increases in hippocampal ERα levels following previous midlife estradiol exposure can have long-lasting impacts on hippocampal function. Activation of ERα can lead to a wide range of changes within a cell, but its actions are often classified into two categories: genomic and nongenomic. The genomic actions of ERα are the classic steroid hormone receptor actions that involve the receptor acting as a transcription factor at estrogen response elements (EREs) to promote genomic changes within a cell (Klinge, 2001). Consistent with the observed impacts on hippocampal memory, we have also shown that previous exposure to estradiol following ovariectomy results in lasting increases in ERE-dependent transcriptional activity in the hippocampi of ovariectomized ERE-luciferase reporter mice (Pollard et al., 2018). Currently, it remains unknown which specific ERE-dependent genes are impacted by previous midlife estradiol exposure that could be involved with the effects of this hormone treatment paradigm on memory. Three potential genes that contain an ERE sequence and are known to regulate memory include *Esr1* (Castles et al., 1997), *Bdnf* (Sohrabji et al., 1995), and *Chat* (Hyder et al., 1999). Whereas *Esr1* is the gene that transcribes ERα, the proteins transcribed by both *Bdnf* and *Chat* are closely associated with the effects of ERα on hippocampal function (Kőszegi et al., 2011; Scharfman and MacLuskey, 2005). Brain-derived neurotrophic factor (BDNF) is involved in hippocampal neurogenesis and neuroprotection in the aging brain (Pencea et al., 2001; Sohrabji and Lewis, 2006). Choline acetyltransferase (ChAT) is the synthesizing enzyme for acetylcholine, a neurotransmitter closely associated with the actions of estrogen in the hippocampus (Gibbs, 1997; Luine, 1985).

In addition to traditional genomic actions associated with nuclear steroid receptors, membrane localized ERα is also able to impact memory in the hippocampus by acting through rapid nongenomic mechanisms. Membrane ERα has been shown to rapidly activate multiple intracellular signaling pathways that influence hippocampal dependent cognition [for review, see (Foster, 2012)]. Rapid effects of estradiol administration on cellular signaling pathways has been implicated in synaptic transmission (Fugger et al., 2001), cell excitability (Kumar and Foster, 2002), NMDA receptor function (Bi et al., 2003), long-term potentiation (Foy et al., 2008), and rapid changes in expression of synaptic proteins including postsynaptic density protein 95 (PSD-95), a protein crucial for stabilizing synaptic changes during long-term potentiation (Akama and McEwen, 2003; Murakami et al., 2015). Because these different actions of ERα are associated with specific locations of the receptor, the subcellular distribution of ERα dictates its function. Currently, it is unknown where the observed increase in ERα following previous midlife estradiol exposure occurs within hippocampal cells.

The overall goal of the current work was to determine mechanisms by which previous exposure to estradiol in midlife, acting through its ability to increase levels of ERα, is able to maintain hippocampal function. To do so we compared impacts of previous estradiol treatment to ongoing estradiol and ovariectomized control treatments on hippocampal gene and protein expression of estrogen-sensitive genes and on the subcellular distribution of hippocampal ERα protein levels. Specifically, we measured gene expression and corresponding protein levels of three estrogen-sensitive genes that contain ERE sequences (*Esr1/*ERα*, Bdnf/*BDNF*, Chat/*ChAT) and one estrogen-sensitive gene without a known ERE sequence but associated with the actions of membrane-bound ERα (*Dlg4/*PSD-95). Subcellular fractionation was performed before measuring ERα protein levels in order to determine the subcellular localization of the receptor in hippocampal cells following previous exposure to estradiol in midlife.

## 2. Materials and Methods

### 2.1 Subjects

Middle-aged female Long-Evans hooded rats, retired breeders (~11 months of age), were purchased from Envigo. Animal care was in accordance with guidelines set by the National Institute of Health Guide for the Care and Use of Laboratory Animals (2011) and the Institutional Animal Care and Use Committees of Tulane University approved all procedures. Rats were housed individually in a temperature-controlled vivarium under a 12-h light, 12-h dark cycle and had unrestricted access to food and water.

### 2.2 Ovariectomy and hormone treatment

Rats were anesthetized by intraperitoneal injections of ketamine (100 mg/kg ip; Bristol Laboratories, Syracuse, NY) and xylazine (7 mg/kg ip; Miles Laboratories, Shawnee, KS) and were ovariectomized. Buprenorphine (0.375 mg/kg; Reckitt Benckiser Health Care) was administered by subcutaneous injection before surgery. Ovariectomy surgery involved bilateral flank incisions through skin and muscle wall and removal of ovaries. Immediately following ovariectomy, rats were implanted with a subcutaneous 5-mm SILASTIC brand capsule (0.058 in. inner diameter and 0.077 in. outer diameter; Dow Corning, Midland, MI) on the dorsal aspect of their necks. Capsules contained either vehicle or 25% 17β-estradiol (Sigma-Aldrich, St. Louis, MO) diluted in vehicle. We have previously shown that implants of these dimensions and estradiol concentrations maintain blood serum estradiol levels in middle-age retired breeders at approximately 37 pg/mL (Bohacek and Daniel, 2007), which falls within physiological range.

### 2.3 Hormone capsule replacement

Forty days after ovariectomy and capsule implantation, rats were anesthetized with ketamine and xylazine and capsules were removed and replaced with a new capsule. Buprenorphine was administered by subcutaneous injection before the start of each surgery. Rats initially receiving vehicle capsules received another vehicle capsule (Vehicle group). Rats initially receiving estradiol capsules either received a new estradiol capsule (Continuous Estradiol group) or instead received a vehicle capsule (Previous Estradiol group).

### 2.4 Euthanasia and tissue collection

Approximately 30 days following capsule replacement surgeries, rats were killed under anesthesia induced by ketamine and xylazine. Hippocampus were dissected and either quick frozen on dry ice for processing for western blotting or placed into tubes containing RNAlater (Qiagen; Hilden, Germany) for RNA extraction and stored at −80°C until further processing.

### 2.5 Hormone treatment verification

Vaginal smears for each rat were collected for at least four consecutive days before capsule replacement in order to confirm hormone treatment for the initial forty-day window. Smears of ovariectomized, cholesterol-treated rats were characterized by a predominance of leukocytes, while smears of ovariectomized, estradiol-treated rats were characterized by a predominance of cornified and nucleated epithelial cells indicating hormone treatment was effective. At the time of euthanasia, a 1-cm sample of the right uterine horn was collected from each rat and weighed to verify hormone treatment during the latter part of the experiment.

### 2.6 Subcellular protein fractionation

Hippocampal tissue from each of the hormone treatment groups (Vehicle, n=10; Continuous Estradiol, n=10; Previous Estradiol, n=10) was lysed in cytosolic extraction buffer and protease inhibitors included in the Sub-Cellular Protein Fractionation Kit for Tissues (Thermo Scientific, Waltham, MA) using the PowerGen 125 handheld homogenizer (Fisher Scientific, San Jose, CA). Homogenate was centrifuged through the Pierce Tissue Strainer at 500 x g for 5 minutes at 4°C. Supernatant containing the cytosolic compartment extract was transferred immediately to a clean tube. The pellet was resuspended by vortexing in membrane extraction buffer containing protease inhibitors then incubated at 4°C for 10 minutes with gentle mixing. Sample was centrifuged at 3000 x g for 5 minutes at 4°C. Supernatant containing the membrane compartment extract was transferred immediately to a clean tube. The pellet was resuspended by vortexing in nuclear extraction buffer and protease inhibitors then incubated at 4°C for 30 minutes with gentle mixing. The sample was then centrifuged at 5000 x g for 5 minutes at 4°C. Supernatant containing the nuclear soluble extract was transferred to a clean tube. The pellet was resuspended by vortexing in chromatin bound extraction buffer containing room temperature nuclear extraction buffer with protease inhibitors, 100mM CaCl2, and Micrococcal Nuclease, then incubated at 37°C for 15 minutes. The sample was then centrifuged at 16,000 x g for 5 minutes at 4°C. Supernatant containing the chromatin bound extract was added to the previously obtained nuclear soluble extract to constitute the nuclear compartment extract. Protein concentration was determined for the cytosolic, membrane, and nuclear fractions of each sample using the Bradford Protein Assay (Thermo Scientific). Each compartment of each sample was diluted 1:1 in Laemlli Sample Buffer (BioRad, Hercules, CA) mixed with 350mM DTT (Sigma-Aldrich) and boiled for 5 minutes. One cytosolic and one membrane sample from the Vehicle group, one membrane and two nuclear samples from the Continuous Estradiol group, and one cytosolic sample from the Previous Estradiol group were excluded from western blotting either due to compartmental contamination or low protein yield.

### 2.7 Subcellular compartment western blotting

For each cytosolic, membrane, and nuclear tissue from each sample, 15ug of protein were loaded onto and separated on a 7.5% TGX SDS-PAGE gel at 250 V for 40 minutes. Molecular weight markers (PageRuler, Thermo Scientific) were included with each run. Proteins were transferred from gels to nitrocellulose membranes at 100 V for 30 minutes. Membranes were blocked with 5% nonfat dry milk in 1% Tween 20/1 Tris-buffered saline (TTBS) with gentle mixing at room temperature for 1 hour. After blocking, membranes were incubated with gentle mixing in primary antibody overnight at 4°C in 1% nonfat dry milk-TTBS. Samples from cytosolic, membrane, and nuclear compartments were incubated with antibodies for ERα (mouse monoclonal, Santa Cruz; 1:750). Samples from cytosolic fractions were incubated with antibodies for cytosolic loading control Enolase (1:2000, Santa Cruz). Samples from membrane fractions were incubated with antibodies for membrane loading control ATP1A1 (1:5000, ProteinTech). Samples from nuclear fractions were incubated with antibodies for nuclear loading control CREB (1:2000, Cell Signaling). Following primary antibody incubation, blots were washed three times for 15 minutes with TTBS. Blots were then incubated with secondary antibodies conjugated to HRP in 5% NFDM-TTBS for one hour at room temperature with gentle mixing. Secondary antibodies used were Goat Anti-Mouse IgG (1:50,000 for ERα, BioRad) and Goat-Anti Rabbit IgG (1:10,000 for enolase, ATP1A1, and CREB; Santa Cruz). Following secondary antibody incubation, bots were washed three times for 15 minutes with TTBS. Blots were then incubated with the chemiluminescent substrate Supersignal West Femto (Fischer Scientific) for 5 minutes and exposed to film (Kodak) for varying times to capture optimal signal intensity. Films were imaged using MCID Core imaging software (InterFocus Imaging Ltd., Cambridge, England), and optical density was measured for bands of interest. Values for each sample represent the percentage of ERα expression relative to the compartment-specific loading control normalized to the mean vehicle values.

### 2.8 RNA extraction

Hippocampal tissue from each of the hormone treatment groups (Vehicle, n=11; Continuous Estradiol, n=11; Previous Estradiol, n=10) was homogenized using the PowerGen 125 handheld homogenizer in QIAzol Lysis Reagent (Qiagen) and extracted using the RNeasy Plus Universal Mini Kit (Qiagen). Briefly, lysate was incubated with chloroform and the aqueous phase was then incubated with ethanol. Sample was then centrifuged and washed in a spin column. RNA was eluted using Rnase-free water. A gDNA eliminator was used to reduce genomic DNA contamination. RNA quality and concentration were determined by gel electrophoresis and UV absorption.

### 2.9 RT-PCR

RNA was quantified using the QuantiFast SYBR Green RT-PCR Kit (Qiagen). Primers used were for *Esr1* (QuantiTect Primer Assays; Qiagen), *Chat*, *Bdnf*, *Dlg4* and the housekeeping gene, *Gapdh* (QuantiTect Primer Assays; Qiagen). All samples were run in triplicate. Reverse transcription was performed at 50°C for 10 min to generate cDNA in a 25 μL reaction volume with 50 μg total RNA. HotStarTaq Plus DNA Polymerase was activated and reverse transcription was ended by 5 min of incubation at 95°C. Following the initial activation step, 40 cycles of 2-step cycling consisting of denaturation for 10 s at 95°C and combined annealing/extension for 30 s at 60°C were repeated. Melting curves were observed to confirm the correct primer usage in each well. Values analyzed represent a mean of triplicates normalized as a percentage of the values for the housekeeping gene, *Gapdh*.

### 2.10 Whole cell tissue processing

Hippocampal tissue from each of the hormone treatment groups (Vehicle, n=8; Continuous Estradiol, n=8; Previous Estradiol, n=10) were homogenized in 10 μl/mg lysis buffer containing 1 mM EGTA, 1 mM EDTA, 20 mM Tris, 1 mM sodium pyrophosphate tetrabasic decahydrate, 4 mM 4-nitrophenyl phosphate disodium salt hexahydrate, 0.1 μM microcystin, and 1% protease inhibitor cocktail (Sigma-Aldrich). Samples were then centrifuged for 15 min at 1000 x g at 4°C. Protein concentration of supernatant determined via Bradford Protein Assay. Each sample diluted 1:1 with Laemmli Sample Buffer (BioRad) mixed with 350 mM DTT, boiled for 10 min, and stored at −80°C until western blotting.

### 2.11 Whole cell western blotting

Twenty-five micrograms of protein were loaded onto a gel and western blotting was performed as described in section 2.7. Primary antibodies used include PSD-95 (Millipore, 1:2000), BCL-2 (Santa Cruz, 1:2000), ChAT (Cell Signalling, 1:2000), BDNF (Santa Cruz, 1:2000), and loading control protein β-actin (Santa Cruz, 1:5000). Secondary antibodies conjugated to HRP were used. Blots were then incubated with the chemiluminescent substrates Supersignal West Femto (PSD-95, BDNF, ChAT) for 5 minutes or ECL Standard (β -actin) for 1 minute and then imaged using the ChemiDocMP Imaging System (BioRad). One sample from the Vehicle group was excluded from analysis for the BDNF western blot due to a transfer bubble over the band of interest for that sample. Optical density x area was measured for bands of interest using MCID Imaging software. Data for each band of interest were normalized to expression of loading control protein β -actin.

### 2.12 Statistical analyses

Data were analyzed by One-Way ANOVA comparing treatment group and subsequent LSD post-hoc testing as appropriate. Researchers were blind to treatment group during western blotting, RT-PCR and data analysis. All data analyses were performed using SPSS software.

## 3. Results

### 3.1 Transcriptional regulation of Esr1 following continuous or previous estradiol exposure in the hippocampus of aging ovariectomized rats

As shown in Figure 1A, there was no effect of hormone treatment on hippocampal expression of *Esr1* (F(2,31)=0.422, p=0.660). These data suggest that elevated hippocampal ERα protein levels following continuous and previous estradiol exposure (Rodgers et al. 2010; Witty et al. 2013) during midlife are not caused by changes in transcription levels in *Esr1*.

**Figure 1.**
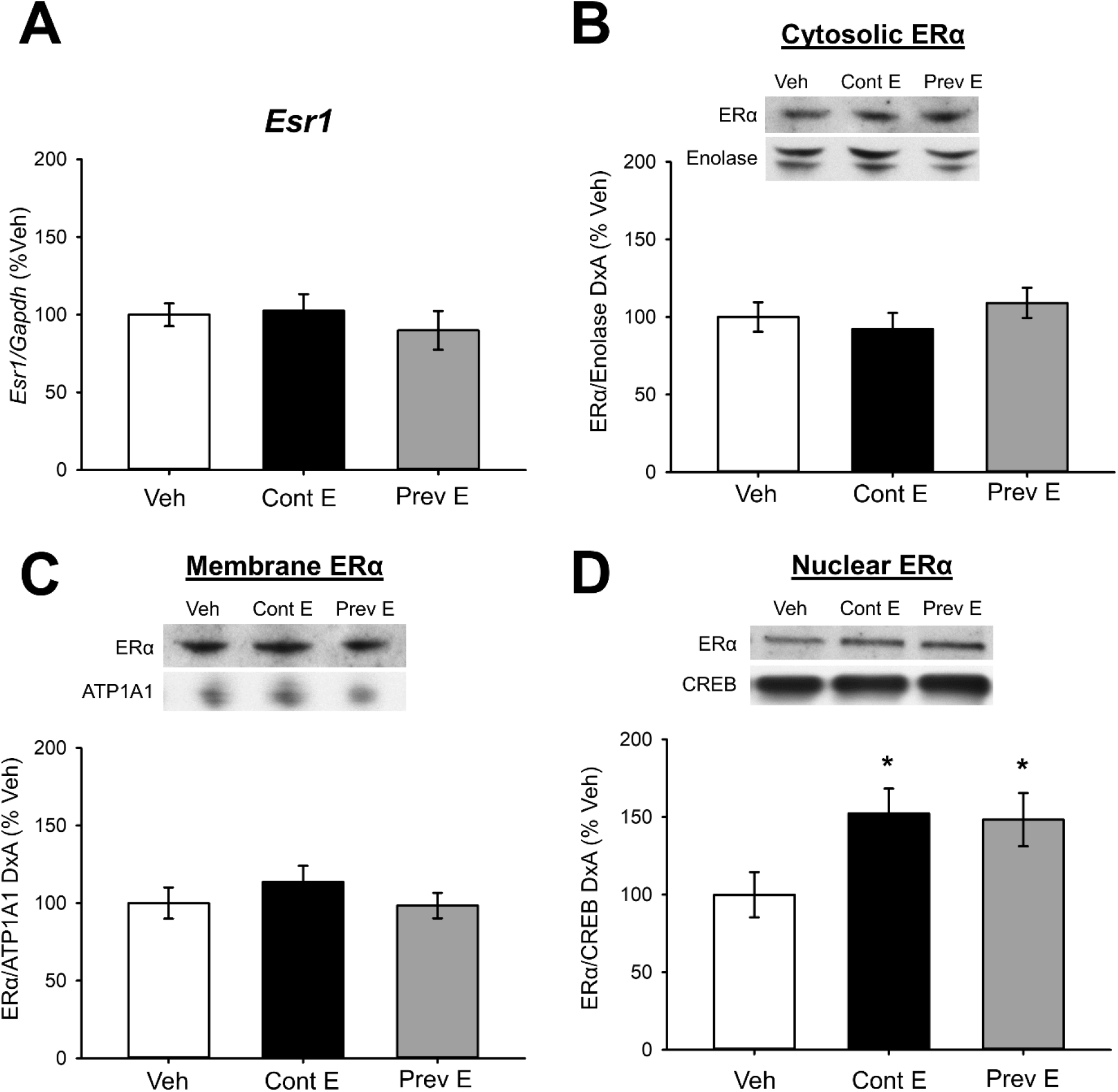
Transcriptional Regulation of Esr1 and subcellular localization of ERα in the hippocampus of ovariectomized rats following continuous or previous exposure to estradiol in midlife. Middle-aged female rats were ovariectomized and treated to one of three hormone conditions via Silastic capsule: Vehicle (Veh), which received a vehicle capsule; Continuous Estradiol (Cont E), which received an estradiol capsule for the duration of the experiment; or Previous Estradiol (Prev E), which received an estradiol capsule for 40 days followed by a vehicle capsule for the duration of the experiment. One month later, hippocampi were processed for either RNA extraction and RT-PCR using primers for *Esr1* and housekeeping gene *Gapdh* or for subcellular fractionation and western blotting for ERα, measured by density x area of ERα/loading control proteins. A) There was no significant effect of hormone treatment on *Esr1* expression in the hippocampus relative to *Gapdh* expression. B-C) There was no significant effect of hormone treatment on cytosolic (B) or membrane (C) ERα. D) There was a significant effect of hormone treatment on levels of nuclear ERα. Post hoc testing revealed increased levels in the Cont E and Prev E groups relative to the Veh group. Data are presented as means ± SEM normalized to percent Vehicle group. *p<.05 vs. Veh

### 3.2 Subcellular localization of ERα in the hippocampus of ovariectomized rats following continuous or previous exposure to estradiol in midlife

Figure 2 displays verification of our subcellular compartment fractionation process of hippocampal tissue. Enolase—an enzyme involved in glycolysis—was used as a cytosolic marker and loading control, appearing predominately in the cytosolic fraction of samples. ATP1A1—a subunit of the sodium potassium pump ATPase—was used as a membrane marker and loading control, appearing predominately in the membrane fraction of samples. The transcription factor cAMP response element-binding protein (CREB) was used as a nuclear marker and loading control, appearing predominately in the nuclear fraction of samples. Verification of compartment fractionation was performed by western blotting for each of the compartment markers using a random selection of samples from each experimental group.

**Figure 2.**
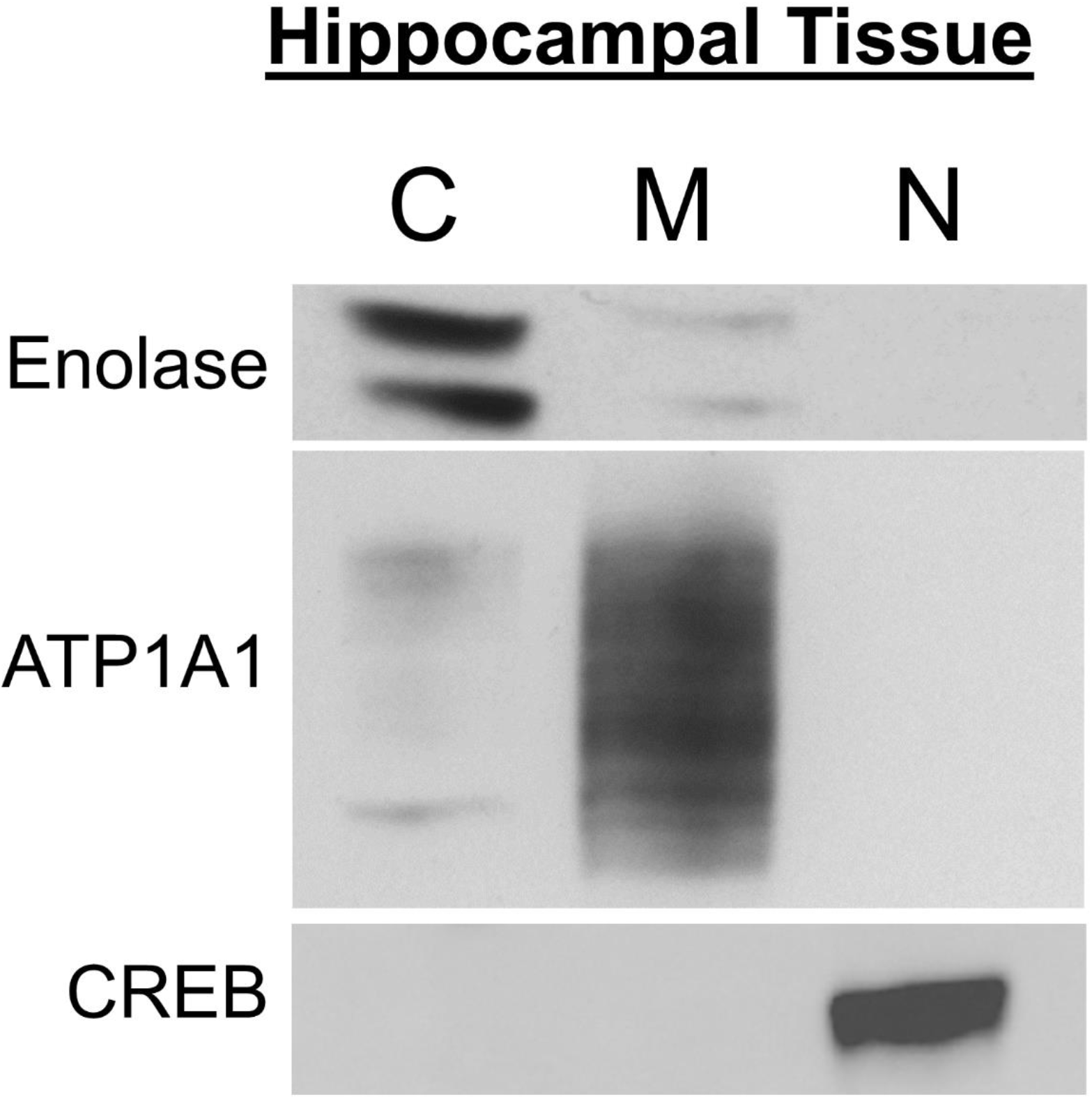
Verification of subcellular compartment fractionation. Hippocampal tissue was processed for subcellular fractionation in order to separate the cytosolic, membrane, and nuclear compartments of cells via consecutive centrifugation steps using a commercially available kit. Compartment separation was verified using western blotting for cytosolic marker enolase, membrane marker ATP1A1, and nuclear marker CREB on samples from all compartments.

As illustrated in Figures 1B-C, there was no effect of hormone treatment on cytosolic (*F*(2,27)=0.727, *p*=0.493) or membrane (*F*(2,27)=0.763, *p*=.477) ERα levels. As illustrated in Figure 1D, there was an effect of hormone treatment on nuclear ERα levels (*F*(2,27)=3.396, *p*=0.0496), with levels increased in both the Continuous Estradiol (*p*=0.033) and Previous Estradiol (*p*=0.036) groups as compared to Vehicle group levels. There was no significant difference in nuclear ERα levels between the Continuous Estradiol and Previous Estradiol groups (*p*=0.863). These results demonstrate that previous exposure to estradiol in midlife results in lasting elevation specifically of nuclear ERα levels in the hippocampus of ovariectomized rats, similar to levels in animals receiving ongoing estradiol treatment.

### 3.3 Hippocampal transcriptional regulation and protein expression of genes that contain ERE sequences following continuous or previous midlife estradiol exposure

As illustrated in Figure 3A, there was an effect of hormone treatment on expression of *Bdnf* (*F*(2,31)=4.180, *p*=0.025), with increased RNA levels in both the Continuous Estradiol (*p*=0.011) and Previous Estradiol (*p*=0.039) groups as compared to the levels in the Vehicle group. No significant difference in RNA levels of *Bdnf* was found between the Continuous Estradiol and Previous Estradiol groups (*p*=0.621). There was an effect of hormone treatment on hippocampal protein levels of BDNF (Figure 3B, *F*(2,24)=6.676, *p*=0.005), with levels in the Previous Estradiol group significantly increased as compared to those in both the Vehicle (*p*=0.023) and the Continuous Estradiol (*p*=0.002). No significant difference in protein levels of BDNF was found between the Vehicle and Continuous Estradiol groups (*p*=0.390).

**Figure 3.**
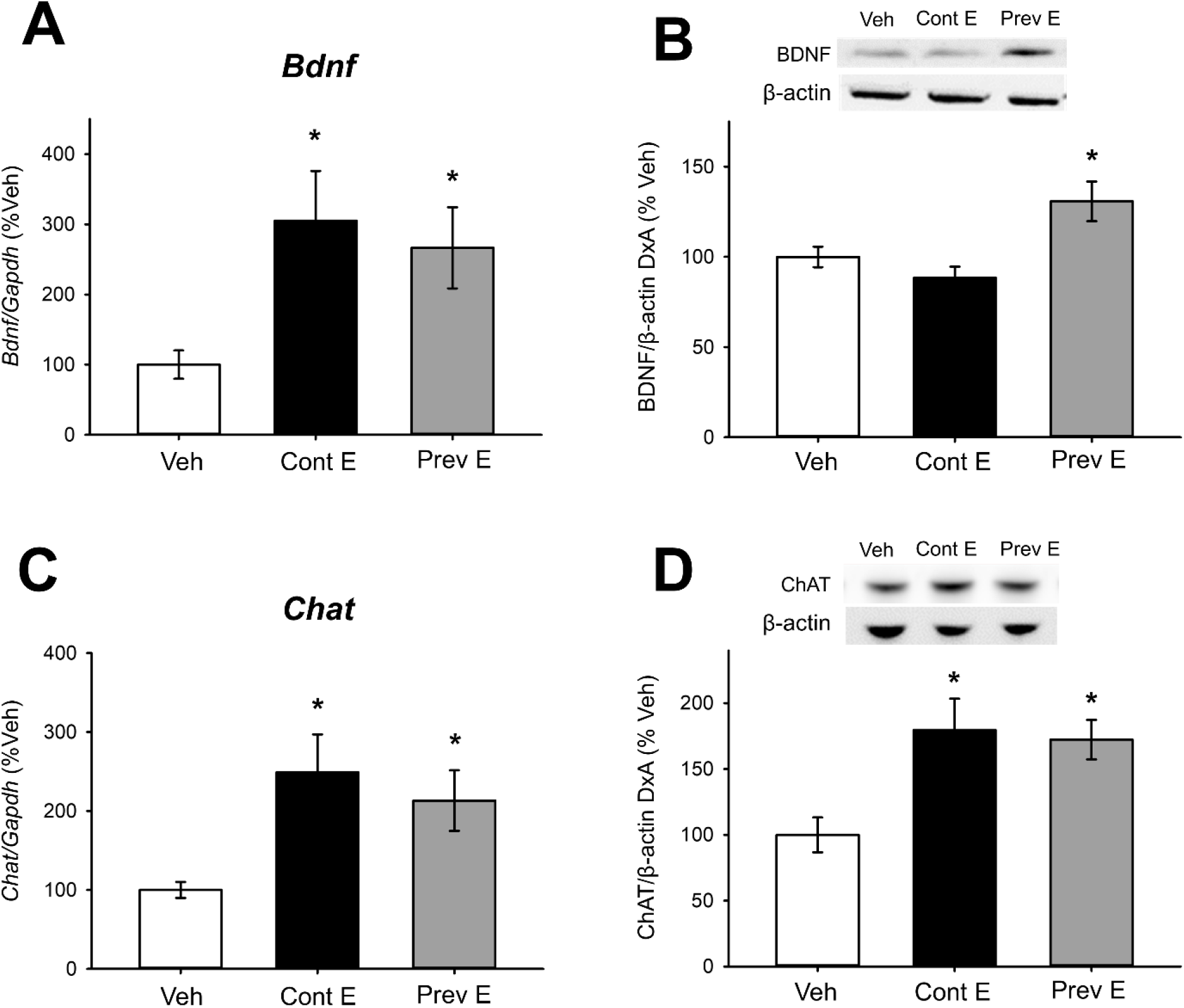
Hippocampal transcriptional regulation and protein expression of genes that contain ERE sequences following continuous or previous midlife estradiol exposure. Middle-aged female rats were ovariectomized and treated to one of three hormone conditions via Silastic capsule: Vehicle (Veh), which received a vehicle capsule; Continuous Estradiol (Cont E), which received an estradiol capsule for the duration of the experiment; or Previous Estradiol (Prev E), which received an estradiol capsule for 40 days followed by a vehicle capsule for the duration of the experiment. One month later, hippocampi were processed for RNA extraction and RT-PCR using primers for *Bdnf, Chat,* and *Gapdh*, or for western blotting for BDNF, ChAT, and β-actin. RT-PCR data were normalized to housekeeping gene *Gapdh* and western blot data were normalized to loading control protein β-actin. A-B) There was a significant effect of hormone treatment on *Bdnf* RNA expression (A) and BDNF protein levels (B) in the hippocampus. Post hoc testing revealed increased *Bdnf* expression in the Cont E and Prev E groups and increased BDNF protein levels in the Prev E group as compared to the Veh group. C-D) There was a significant effect of hormone treatment on *ChAT* RNA expression (C) and ChAT protein levels (D) in the hippocampus. Post hoc testing revealed increased *Chat* expression and increased ChAT protein levels in the Cont E and Prev E groups as compared to the Veh group. *p<.05 vs. Veh

As illustrated in Figure 3C, there was an effect of hormone treatment on expression of *Chat* (Figure 2C, *F*(2,31)=4.810, *p*=0.016), with increased RNA levels in both the Continuous Estradiol (*p*=0.006) and Previous Estradiol (*p*=0.035) groups as compared to the levels in the Vehicle group. No significant difference in RNA levels of *Chat* was found between the Continuous Estradiol and Previous Estradiol groups (*p*=0.492). There was an effect of hormone treatment on hippocampal protein levels of ChAT (Figure 3D, *F*(2,25)=5.944, *p*=0.008), with levels in the Continuous Estradiol (*p*=0.005) and Previous Estradiol (*p*=0.007) groups significantly increased as compared to those in the Vehicle group. No significant difference in ChAT protein levels of was found between the Continuous Estradiol and Previous Estradiol groups (*p*=0.770).

### 3.4 Hippocampal transcriptional regulation and protein expression of one gene that does not contain an ERE sequence following continuous or previous midlife estradiol exposure

As illustrated in Figure 4A, there was an effect of hormone treatment on expression of *Dlg4* (*F*(2,31)=3.812, *p*=0.034), with increased RNA levels in both the Continuous Estradiol (*p*=0.034) and Previous Estradiol (*p*=0.018) groups as compared to the levels in the Vehicle group. No significant difference in RNA levels of *Dlg4* was found between the Continuous Estradiol and Previous Estradiol groups (*p*=0.731). There was no effect of hormone treatment on hippocampal protein levels of PSD-95 (*F*(2,25)=0.088, *p*=0.916).

**Figure 4.**
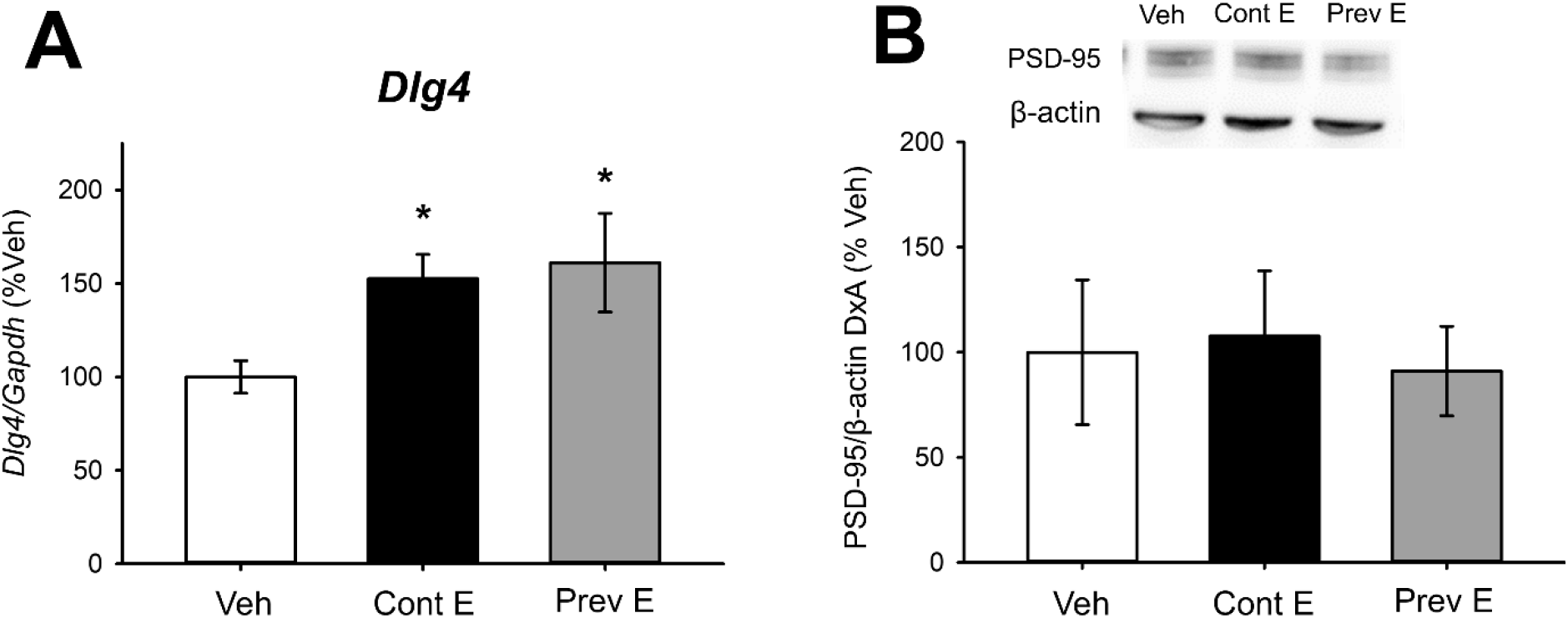
Hippocampal transcriptional regulation and protein expression of gene that does not contain an ERE sequence following continuous or previous midlife estradiol exposure. Middle-aged female rats were ovariectomized and treated to one of three hormone conditions via Silastic capsule: Vehicle (Veh), which received a vehicle capsule; Continuous Estradiol (Cont E), which received an estradiol capsule for the duration of the experiment; or Previous Estradiol (Prev E), which received an estradiol capsule for 40 days followed by a vehicle capsule for the duration of the experiment. One month later, hippocampi were processed for RNA extraction and RT-PCR using primers for *Dlg4* and *Gapdh*, or for western blotting for PSD-95 and β-actin. RT-PCR data were normalized to housekeeping gene *Gapdh* and western blot data were normalized to loading control protein β-actin. A) There was a significant effect of hormone treatment on *Dlg4* RNA expression in the hippocampus. Post hoc testing revealed increased *Dlg4* expression in the Cont E and Prev E groups as compared to the Veh group. B) There was no effect of hormone treatment on PSD-95 protein levels. *p<.05 vs. Veh

## 4. Discussion

Results of the present studies reveal that previous exposure to estradiol during midlife has lasting impacts on hippocampal function through sustained transcriptional activity of ERα that persists long after estradiol treatment has ended. Previous estradiol exposure resulted in lasting increases in the nuclear pool of hippocampal ERα as well as long-term upregulation of BDNF and ChAT in aging ovariectomized rats—effects that were similar to those observed in ovariectomized animals with ongoing estradiol exposure. Specifically, we showed that both continuous and previous estradiol increased nuclear ERα protein levels in the hippocampus of aging, ovariectomized rats one month after cessation of estradiol treatment. Animals from the Previous Estradiol group also had elevated hippocampal expression of two ERE-dependent genes (*Bdnf, Chat*) and their corresponding proteins (BDNF, ChAT), as well as elevated expression of one non-ERE-dependent gene (*Dlg4*), as compared to animals from the Vehicle group. Animals receiving ongoing estradiol exposure displayed similar, but not identical, expression patterns in the hippocampus, with elevated gene expression of *Bdnf*, *Chat*, and *Dlg4*, but only elevated protein levels of ChAT as compared to animals from the Vehicle group. Overall, these findings indicate a crucial role for ERα in maintaining hippocampal memory in ovariectomized animals through sustained transcriptional activity of the receptor following previous estradiol exposure in midlife in a pattern that is comparable to that observed in the presence of circulating estradiol.

Maintained hippocampal protein levels of ERα following continuous and previous estradiol exposure in ovariectomized rats has been associated with enhanced hippocampal-dependent memory (Rodgers et al., 2010). As a nuclear steroid hormone receptor, ERα can act in a variety of functions within a cell. Here we demonstrated through subcellular compartment fractionation that ERα is specifically increased in the nuclear compartment of hippocampal cells following continuous or previous midlife estradiol exposure in ovariectomized rats, although the receptor is also present in the cytosol and membrane compartments in all hormone treatment groups. This finding aligns with recent work from our lab demonstrating that short-term estradiol treatment immediately after ovariectomy results in lasting enhancements in hippocampal-memory and increased ERE-dependent transcriptional activity in mice (Pollard et al., 2018). Together, it strengthens the connection between maintained protein levels of ERα following previous exposure to estradiol in midlife and enhanced memory via sustained transcriptional activity of ERα in the hippocampus.

The current results, in which we did not see estradiol-induced impacts on *Esr1* transcription levels, suggest that mechanisms by which previous estradiol treatment maintains levels of ERα in the hippocampus long after the termination of the treatment does not involve transcriptional regulation. Besides modification of ERα gene transcription, ERα levels can be impacted by changes in receptor degradation rate. ERα is degraded via the ubiquitin-proteasomal pathway (Tateishi et al., 2004). When the receptor is unliganded, the E3 ubiquitin ligase, C terminus of Hsc70-interacting protein (CHIP) binds ERα and targets it for ubiquitination and proteasomal degradation (Fan et al., 2005). Long-term estrogen deprivation following OVX increases interaction between CHIP and ERα and thus subsequent ubiquitination of ERα (Zhang et al., 2011). Prior results from our lab indicate that previous estradiol treatment may prevent effects of estrogen deprivation on ERα degradation. The same 40 days of previous estradiol treatment used in the current study resulted in lasting decreased association between ERα and CHIP in parallel to increased protein levels of ERα when measured one month following termination of the previous estradiol treatment (Black et al., 2016). Together, these results with those of the current work provide support for the hypothesis that lasting changes in levels of ERα resulting from previous exposure to midlife estradiol are primarily due to lasting changes in protein degradation rather than transcription.

The ability of estradiol treatment to increase levels of ERα specifically in the nucleus is consistent with its ability to impact estradiol-sensitive genes and proteins. We found impacts of both continuous and previous estradiol treatments on hippocampal gene expression and protein levels in two ERE-dependent genes known to impact memory. Animals that had previously been exposed to estradiol following ovariectomy displayed upregulated expression of *Bdnf* and *Chat* in the hippocampus one month after termination of estradiol exposure. These increases in gene expression observed in the Previous Estradiol group were comparable to those found in the hippocampi of animals receiving ongoing estradiol exposure, demonstrating that short-term estradiol exposure during midlife can have lasting effects on hippocampal gene expression. There were, however, differences between the Continuous Estradiol and Previous Estradiol group in the expression patterns of the proteins associated with these genes.

Significant increases in hippocampal proteins levels of both BDNF and ChAT were found in the Previous Estradiol group, consistent with the observed increases in gene expression of *Bdnf* and *Chat* described above. These findings were also consistent with earlier findings from our lab that previous estradiol exposure during midlife results in lasting increases in levels of ChAT in the hippocampus (Rodgers et al., 2010; Witty et al., 2013) but which has not been shown before for hippocampal BDNF levels. In contrast to the effects observed in the Previous Estradiol group, animals in the Continuous Estradiol group showed significantly increased hippocampal protein levels of ChAT, but not BDNF, despite showing increased gene expression for both *Bdnf* and *Chat*. Several earlier studies have demonstrated that estradiol exposure increases *Chat* mRNA (Gibbs, 1996), elevates ChAT protein expression (Gibbs, 1997; Rodgers et al., 2010) and increases acetylcholine release in the hippocampus of ovariectomized rodents (Gabor et al., 2003; Gibbs, 1997). However, the relationship between estradiol exposure, *Bdnf* mRNA, and BDNF protein levels in ovariectomized animals is less clear. Consistent with our findings, several—but not all (Cavus and Duman, 2003)—studies found continuous estradiol treatment to increase hippocampal mRNA levels of *Bdnf* in ovariectomized animals (Berchtold et al., 2001; Liu et al., 2001; Singh et al., 2005). There is however no clear consensus as to the effect of estradiol treatment on hippocampal BDNF protein levels, with some studies showing increased protein expression throughout the hippocampus (Berchtold et al., 2001), some showing increased BDNF levels only in specific subregions of the hippocampus (Zhou et al., 2005), and others showing decreased hippocampal protein levels in ovariectomized animals treated with estradiol (Gibbs, 1999). The results of the current studies therefore add to this complex story by demonstrating that previous exposure to estradiol during midlife results in lasting increases in hippocampal mRNA and protein expression of BDNF, whereas continuous exposure to estradiol only results in increased mRNA levels. Nevertheless, both hormone treatments result in lasting increases in mRNA and protein levels of ChAT, another ERE-dependent gene critical for memory. As described above regarding estrogenic regulation of *Esr1* mRNA, the relationship between estrogen receptor activity, mRNA expression, and protein levels can vary greatly due to brain region, age, and duration of estrogen exposure, among other variables (Scharfman and MacLuskey, 2005).

Finally, our findings demonstrate that continuous or previous estradiol exposure following ovariectomy can also result in increased transcription of non-ERE-dependent genes, with both the Continuous Estradiol and Previous Estradiol groups showing increased hippocampal expression of *Dlg4* as compared to the Vehicle group. Interestingly, there was no observed change in levels of PSD-95, the protein transcribed by *Dlg4*, in either hormone treatment group. Estradiol treatment has repeatedly been shown to increase protein levels of PSD-95 in the hippocampus (Nelson et al., 2014; Waters et al., 2009), a rapid effect that is attributed to the PI3K-Akt signaling pathway activated by membrane ERα (Akama and McEwen, 2003; Murakami et al., 2015). Because we observed no change in membrane levels of ERα following continuous or previous estradiol treatments, it is unclear how increased ERα in the nucleus of hippocampal cells can result in lasting changes in gene expression but not protein levels of PSD-95. Future studies should investigate potential crosstalk between membrane and nuclear ERα and its impact on synaptic proteins following previous midlife estradiol exposure.

Together, the results of the present study indicate a critical role for ERα as a transcriptional regulator of hippocampal function in both the presence and absence of circulating estrogens. Previous exposure to estradiol in midlife results in lasting increases in nuclear ERα activity in the hippocampus, resulting in increased transcription of genes important for hippocampal function and enhanced hippocampal dependent memory (Rodgers et al., 2010). Future work should thoroughly examine the mechanisms through which ERα can influence hippocampal function in the absence of circulating estrogens, though previous work from our lab indicates a role for brain-derived neuroestrogens (Baumgartner et al., 2019) as well as ligand-independent activation of ERα by growth factors including insulin-like growth factor-1 (IGF-1) (Grissom and Daniel, 2016; Witty et al., 2013).

Ultimately these findings have important implications for women who use short-term estrogen treatment to treat their menopause symptoms. Several studies, including those of women who underwent surgical menopause earlier in life than natural menopause (Bove et al., 2014; Rocca et al., 2014), suggest a lasting benefit for cognition of short-term estrogen use immediately following menopause (Bagger et al., 2005; Whitmer et al., 2011). Ongoing large-scale clinical studies such as the Kronos Early Estrogen Prevention Study (KEEPS) will provide more insight into the long-term impacts of short-term estrogen use in midlife (Wharton et al., 2013). The results of the current study suggest a potential mechanism for short-term midlife estrogen use to have lasting impacts on cognition by maintaining transcriptional activity of nuclear ERα in the hippocampus. Furthermore, these findings emphasize the critical impact that ERα can have on the aging female brain in the absence of circulating estrogens.

## Notes

Funding: This work was supported by the National Institute on Aging under Grant RF1AG041374.

### Competing Interest Statement

The authors have declared no competing interest.

## References

Akama, K., McEwen, B., 2003. Estrogen stimulates postsynaptic density-95 rapid protein synthesis via the Akt/protein kinase B pathway. J Neurosci 23(6), 2333–2339.

Bagger, Y., Tankó, L., Alexandersen, P., Qin, G., Christiansen, C., 2005. Early postmenopausal hormone therapy may prevent cognitive impairment later in life. Menopause 12(1), 12–17.

Baumgartner, N., Grissom, E., Pollard, K., McQuillen, S., Daniel, J., 2019. Neuroestrogen-Dependent Transcriptional Activity in the Brains of ERE-Luciferase Reporter Mice following Short- and Long-Term Ovariectomy. eNeuro 6(5), ENEURO.0275-0219.2019.

Baxter, M., Santistevan, A., Bliss-Moreau, E., Morrison, J., 2018. Timing of cyclic estradiol treatment differentially affects cognition in aged female rhesus monkeys. Beh Neurosci 132(4), 213–223.

Berchtold, N., Kesslak, J., Pike, C., Adlard, P., Cotman, C., 2001. Estrogen and exercise interact to regulate brain-derived neurotrophic factor mRNA and protein expression in the hippocampus. European Journal of Neuroscience 14(12), 1992–2002.

Bi, R., Foy, M., Thompson, R., Baudry, M., 2003. Effects of estrogen, age, and calpain on MAP kinase and NMDA receptors in female rat brain. Neurobiol Aging 24, 977–983.

Black, K.L., Baumgartner, N.E., Daniel, J.M., 2018. Lasting impact on memory of midlife exposure to exogenous and endogenous estrogens. Behav. Neurosci. 132(6), 547–551.

Black, K.L., Witty, C.F., Daniel, J.M., 2016. Previous Midlife Oestradiol Treatment Results in Long-Term Maintenance of Hippocampal Oestrogen Receptor alpha Levels in Ovariectomised Rats: Mechanisms and Implications for Memory. J Neuroendocrinol 28(10).

Bohacek, J., Daniel, J.M., 2007. Increased daily handling of ovariectomized rats enhances performance on a radial-maze task and obscures effects of estradiol replacement. Horm Behav 52(2), 237–243.

Bove, R., Secor, E., Chibnik, L., Barnes, L., Schneider, J., Bennett, D., De Jager, P., 2014. Age at surgical menopause influences cognitive decline and Alzheimer pathology in older women. Neurology 82(3), 222–229.

Castles, C., Oesterreich, S., Hansen, R., Fugua, S., 1997. Auto-regulation of the estrogen receptor promoter. Ster Biochem Mol Bio 62(2-3), 155–163.

Cavus, I., Duman, R., 2003. Influence of estradiol, stress, and 5-HT2A agonist treatment on brain-derived neurotrophic factor expression in female rats. Biol. Psychiatry 54(1), 59–69.

Chen, W., Manson, J., Hankinson, S., Rosner, B., Holmes, M., Willett, W., Colditz, G., 2006. Unopposed estrogen therapy and the risk of invasive breast cancer. Arch Intern Med 166, 1027–1032.

Daniel, J.M., Witty, C.F., Rodgers, S.P., 2015. Long-term consequences of estrogens administered in midlife on female cognitive aging. Horm Behav 74, 77–85.

Fan, M., Park, A., Nephew, K., 2005. CHIP (carboxyl terminus of Hsc70-interacting protein) promotes basal and geldanamycin-induced degradation of estrogen receptor-alpha. Mol Endocrinol 19(12), 2901–2914.

Foster, T., 2012. Role of Estrogen Receptor Alpha and Beta Expression and Signaling on Cognitive Function During Aging. Hippocampus 22(4), 656–669.

Foy, M., Baudry, M., Foy, J., Thompson, R., 2008. 17beta-estradiol modifies stress-induced and age-related changes in hippocampal synaptic plasticity. Behav Neurosci 122(2), 301–309.

Fugger, H., Kumar, A., Lubahn, D., Korach, K., Foster, T., 2001. Examination of estradiol effects on the rapid estradiol mediated increase in hippocampal synaptic transmission in estrogen receptor alpha knockout mice. Neurosci Lett 309(3), 207–209.

Gabor, R., Nagle, R., Johnson, D., Gibbs, R., 2003. Estrogen enhances potassium-stimulated acetylcholine release in the rat hippocampus. Brain Res 962(1), 244–247.

Gibbs, R., 1996. Fluctuations in relative levels of choline acetyltransferase mRNA in different regions of the rat basal forebrain across the estrous cycle: effects of estrogen and progesterone. J Neuro 16(3), 1049–1055.

Gibbs, R., 1997. Effects of estrogen on basal forebrain cholinergic neurons vary as a function of dose and duration of treatment. Brain Research 757(1), 10–16.

Gibbs, R., 1999. Treatment with estrogen and progesterone affects relative levels of brain-derived neurotrophic factor mRNA and protein in different regions of the adult rat brain. Brain Research 844(1-2), 20–27.

Grissom, E.M., Daniel, J.M., 2016. Evidence for Ligand-Independent Activation of Hippocampal Estrogen Receptor-alpha by IGF-1 in Hippocampus of Ovariectomized Rats. Endocrinology 157(8), 3149–3156.

Hyder, S., Chiappetta, C., Stancel, G., 1999. Interaction of human estrogen receptors alpha and beta with the same naturally occurring estrogen response elements. Biochem Pharmacol 57(6), 597–601.

Klinge, C., 2001. Estrogen receptor interaction with estrogen response elements. Nucleic Acids Res 29(14), 2905–2919.

Koebele, S.V., Bimonte-Nelson, H.A., 2017. The endocrine-brain-aging triad where many paths meet: female reproductive hormone changes at midlife and their influence on circuits important for learning and memory. Exp Gerontol 94, 14–23.

Kőszegi, Z., Szego, É., Cheong, R., Tolod-Kemp, E., Ábrahám, I., 2011. Postlesion Estradiol Treatment Increases Cortical Cholinergic Innervations via Estrogen Receptor-α Dependent Nonclassical Estrogen Signaling *in Vivo*. Endocrinology 152(9), 3471–3482.

Kumar, A., Foster, T., 2002. 17beta-estradiol benzoate decreases the AHP amplitude in CA1 pyramidal neurons. J Neurophysiol 88(2), 621–626.

Liu, Y., Fowler, C., Young, L., Yan, Q., Insel, T., Wang, Z., 2001. Expression and estrogen regulation of brain-derived neurotrophic factor gene and protein in the forebrain of female prairie voles. J Comp Neuro 433(4), 499–514.

Luine, V., 1985. Estradiol increases choline acetyltransferase activity in specific basal forebrain nuclei and projection areas of female rats. Exp Neurol 89(2), 484–490.

Luine, V., Frankfurt, M., 2020. Estrogenic regulation of memory: The first 50 years. Hormones and Behavior 121, 104711.

Ma, S., Tang, N., Leung, G., Fung, A., Lam, L., 2014. Estrogen receptor α polymorphisms and the risk of cognitive decline: A 2-year follow-up study. Am J Geriatr Psychiatry. 22(5), 489–498.

Ma, S., Tang, N., Tam, C., Lui, V., Lau, E., Zhang, Y., Chiu, H., Lam, L., 2009. Polymorphisms of the estrogen receptor alpha (ESR1) gene and the risk of Alzheimer’s disease in a southern Chinese community. Int Psychogeriatr. 21(5), 977–986.

Maki, P., Dennerstein, L., Clark, M., Guthrie, J., LaMontagne, P., Fornelli, D., Little, D., Henderson, V., Resnick, S., 2011. Perimenopausal use of hormone therapy is associated with enhanced memory and hippocampal function later in life. Brain Research 1379, 232–243.

Murakami, G., Hojo, Y., Ogiue-Ikeda, M., Mukai, H., Chambon, P., Nakajima, K., Ooishi, Y., Kimoto, T., Kawato, S., 2015. Estrogen receptor KO mice study on rapid modulation of spines and long-term depression in the hippocampus. Brain Res 1621, 133–146.

Nelson, B., Springer, R., Daniel, J., 2014. Antagonism of brain insulin-like growth factor-1 receptors blocks estradiol effects on memory and levels of hippocampal synaptic proteins in ovariectomized rats. Psychopharmacology (Berl) 231(5), 899–907.

Pencea, V., Bingaman, K., Wiegand, S., Luskin, M., 2001. Infusion of brain-derived neurotrophic factor into the lateral ventricle of the adult rat leads to new neurons in the parenchyma of the striatum, septum, thalamus, and hypothalamus. J Neurosci 21(17), 6706–6717.

Pollard, K.J., Wartman, H.D., Daniel, J.M., 2018. Previous estradiol treatment in ovariectomized mice provides lasting enhancement of memory and brain estrogen receptor activity. Horm Behav 102, 76–84.

Rocca, W., Grossardt, B., Shuster, L., 2014. Oophorectomy, estrogen, and dementia: a 2014 update. Mol Cell Endocrinol 389(1-2), 7–12.

Rodgers, S.P., Bohacek, J., Daniel, J.M., 2010. Transient estradiol exposure during middle age in ovariectomized rats exerts lasting effects on cognitive function and the hippocampus. Endocrinology 151(3), 1194–1203.

Ryan, J., Carrière, I., Carcaillon, L., Dartigues, J., Auriacombe, S., Rouaud, O., Berr, C., Ritchie, K., Scarabin, P., Ancelin, M., 2014. Estrogen receptor polymorphisms and incident dementia: the prospective 3C study. Alzheimers Dement 10(1), 27–35.

Santen, R., Allred, D., Ardoin, S., Archer, D., Boyd, N., Braunstein, G., Burger, H., Colditz, G., Davis, S., Gambacciani, M., Gower, B., Henderson, V., Jarjour, W., Karas, R., Kleerekoper, M., Lobo, R., Manson, J., Marsden, J., Martin, K., Martin, L., Pinkerton, J., Rubinow, D., Teede, H., Thiboutot, D., Utian, W., 2010. Postmenopausal hormone therapy: An endocrine society scientific statement. Clin Endocrinol Metab 95, s1–s66.

Scharfman, H., MacLuskey, N., 2005. Similarities between actions of estrogen and BDNF in the hippocampus: coincidence or clue? Trends in Neurosci 28(2).

Singh, M., Meyer, E., Simpkins, J., 2005. The effect of ovariectomy and estradiol replacement on brain-derived neurotrophic factor messenger ribonucleic acid expression in cortical and hippocampal brain regions of female Sprague-Dawley rats. Endocrinology 136(5).

Sohrabji, F., Lewis, D., 2006. Estrogen-BDNF interactions: implications for neurodegenerative diseases. Front Neuroendocrinol 27(4), 404–414.

Sohrabji, F., Miranda, R., Toran-Allerand, C., 1995. Identification of a putative estrogen response element in the gene encoding brain-derived neurotrophic factor. Proc Nat Acad Sci USA 92(24), 11110–11114.

Tateishi, Y., Kawabe, Y., Chiba, T., Murata, S., Ichikawa, K., Murayama, A., Tanaka, K., Baba, T., Kato, S., Yanagisawa, J., 2004. Ligand-dependent switching of ubiquitin-proteasome pathways for estrogen receptor. EMBO J 23(24), 4813–4823.

Waters, E., Mitterling, K., Jl, S., Mazid, S., McEwen, B., Milner, T., 2009. Estrogen receptor alpha and beta specific agonists regulate expression of synaptic proteins in rat hippocampus. Brain Res 1290, 1–11.

Wharton, W., Gleason, C., Miller, V., Asthana, S., 2013. Rationale and design of the Kronos Early Estrogen Prevention Study (KEEPS) and the KEEPS Cognitive and Affective sub study (KEEPS Cog). Brain Res 1514, 12–17.

Whitmer, R., Quesenberry, C., Zhou, J., Yaffe, K., 2011. Timing of hormone therapy and dementia: the critical window theory revisited. Ann Neurol 69(1), 163–169.

Witty, C., Foster, T., Semple-Rowland, S., Daniel, J., 2012. Increasing hippocampal estrogen receptor alpha levels via viral vectors increases MAP kinase activation and enhances memory in aging rats in the absence of ovarian estrogens. PLoS One 7(12), e51385.

Witty, C., Gardella, L., Perez, M., Daniel, J., 2013. Short-term estradiol administration in aging ovariectomized rats provides lasting benefits for memory and the hippocampus: a role for insulin-like growth factor-I. Endocrinology 154(2), 842–852.

Yaffe, K., Lindquist, K., Sen, S., Cauley, J., Ferrell, R., Penninx, B., Harris, T., Li, R., Cummings, S., 2009. Estrogen receptor genotype and risk of cognitive impairment in elders: findings from the Health ABC study. Neurobiol Aging 30(4), 607–614.

Zhang, Q.G., Han, D., Wang, R.M., Dong, Y., Yang, F., Vadlamudi, R.K., Brann, D.W., 2011. C terminus of Hsc70-interacting protein (CHIP)-mediated degradation of hippocampal estrogen receptor-alpha and the critical period hypothesis of estrogen neuroprotection. Proc Natl Acad Sci USA 108(35), E617–624.

Zhou, J., Zhang, H., Cohen, R., Pandey, S., 2005. Effects of estrogen treatment on expression of brain-derived neurotrophic factor and cAMP response element-binding protein expression and phosphorylation in rat amygdaloid and hippocampal structures. Neuroendocrinology 81(5), 294–310.

